# Identification of new transmembrane proteins concentrated at the nuclear envelope using organellar proteomics of mesenchymal cells

**DOI:** 10.1101/486415

**Authors:** Li-Chun Cheng, Sabyasachi Baboo, Cory Lindsay, Liza Brusman, Salvador Martinez-Bartolomé, Olga Tapia, Xi Zhang, John R. Yates, Larry Gerace

## Abstract

The nuclear envelope (NE) is an endoplasmic reticulum (ER) subdomain that contains characteristic components dedicated to nuclear functions. These include nuclear pore complexes (NPCs) – the channels for nucleocytoplasmic transport, and the nuclear lamina (NL) – a scaffold for NE and chromatin organization at the nuclear periphery. Since numerous human diseases associated with NE/NL proteins occur in mesenchyme-derived cells, a more comprehensive characterization of proteins concentrated at the NE in these cell types is warranted. Accordingly, we used proteomics to analyze NE and other subcellular fractions isolated from mesenchymal stem cells and from differentiated adipocytes and myocytes. We evaluated the proteomics datasets to calculate relative protein enrichment in the NE fraction, using a spectral abundance-based scoring system that accurately described most benchmark proteins. We then examined five high-scoring transmembrane proteins expressed in all three cell types that were not previously known to be enriched at the NE. Using quantitative immunofluorescence microscopy to track ectopically expressed proteins, we validated that all five of these components are substantially concentrated at the NE of multiple cell types. One (Itprip) is exposed to the outer nuclear membrane, a second (Smpd4) is enriched at the NPC, and the three others (Mfsd10, Tmx4, and Arl6ip6) are suggested to reside in the inner nuclear membrane. Considering their sequences and other features, these proteins provide new focal points for studying the functions and membrane dynamics of the NE. Our datasets should be useful for identifying additional NE-concentrated proteins, and for evaluating candidates that are identified in screening.

## Introduction

The nuclear envelope (NE), which forms the membrane boundary of the nucleus, segregates the genome and chromosome-associated metabolism from the cytoplasm. It is a specialized endoplasmic reticulum (ER) sub-domain containing an outer nuclear membrane (ONM) that has continuity and functional similarity with the peripheral ER, and an inner nuclear membrane (INM) with distinctive properties [1,2]. The lipid bilayers of the ONM and INM are joined at nuclear pore complexes (NPCs), massive supramolecular protein assemblies that provide passageways for molecular transport across the NE [3,4]. The NPC is formed from multiple copies of ~30 polypeptides (termed nucleoporins or Nups). A subset of Nups provide scaffolding for the NPC, whereas others, particularly those containing Phe-Gly repeats (termed “FG Nups”), form a diffusion barrier across the NPC and provide binding sites for nuclear transport receptors [3-5].

In higher eukaryotic cells, the INM is lined by the nuclear lamina (NL) – a protein scaffold whose backbone contains a polymer of nuclear lamins, type V intermediate filament proteins [6,7]. Three major lamin subtypes are expressed in the majority of mammalian cells: lamins A/C, B1 and B2 [6,7]. In addition to lamins, at least 20 widely expressed polypeptides are concentrated at the INM [1,6,7]. The NL has been implicated in nuclear structure and mechanics, tethering of heterochromatin and the cytoplasmic cytoskeleton to the NE, and regulation of signaling and gene expression [1,6,7]. Consistent with this wide array of functions, mutations in the genes for lamins and associated proteins have been found to cause a spectrum of human diseases (termed “laminopathies”) [8,9]. Many of these diseases target specific tissues, commonly of mesenchymal origin.

Most of the known INM-enriched proteins have one or more transmembrane (TM) segments [1,10]. Following insertion in the peripheral ER and ONM, these TM proteins are thought to become concentrated at the INM by lateral diffusion in the lipid bilayer around the NPC, in conjunction with binding to the NL and/or other intranuclear components [10, 11]. Movement is bi-directional, and the degree to which specific NE proteins are localized to the INM vs peripheral ER can vary with different cell types or physiological states [12, 13].

Consistent with this diffusion-retention model, ~1/3 of transmembrane proteins in the yeast genome have the ability to reach the INM, even though most do not appear to be concentrated there or have nucleus-specific functions [14]. The principles of the diffusion-retention model also appear to specify NE localization of the LINC complex, an interconnected assembly of TM proteins spanning the INM (SUN-domain proteins) and ONM (nesprins) that is responsible for attaching cytoplasmic cytoskeletal filaments to the NL [15,16].

The detailed protein composition of the NL/INM in mammals remains incompletely understood, and it is likely that low-abundance proteins and/or those with cell type-selective expression patterns remain to be revealed. Proteomics analysis has identified numerous TM proteins in isolated NE fractions of different cell types [17-19], but it remains unclear whether most of these proteins are concentrated at the NE relative to the peripheral ER, or are more general ER residents that by default can diffuse into the contiguous nuclear membranes. An important goal that remains is a comprehensive characterization of proteins that are concentrated at the NE relative to the peripheral ER, as these proteins *de facto* are likely to have specific functions for the nucleus.

In this study, we used proteomics to characterize NEs and other subcellular fractions isolated from cultured mesenchymal stem cells (MSCs), and from correspondingly differentiated adipocytes and myocytes. We implemented a scoring system with the datasets to describe the relative enrichment of individual proteins in the NE fraction. This system accurately represented most of the TM proteins known to be concentrated at the NE, supporting its use to predict new candidates. We selected five of the high-scoring new candidates expressed in all three mesenchymal cell types for direct evaluation by quantitative immunofluorescence microscopy. Our results revealed that all of these are substantially concentrated at the NE: one is enriched at the NPC, one occurs in the ONM, and the remainder appear to be localized to the INM. The sequence homologies and other features of these proteins indicate that they are new windows for understanding the functions and dynamics of the NE. Our datasets provide a resource for evaluating the potential NE localization of membrane proteins detected in proteomics and other screens, and should facilitate the identification of additional NE-concentrated proteins.

## Results

The frequent manifestation of laminopathies in cells of mesenchymal origin [8,9] prompted us to carry out NE proteomics on the murine C3H10T1/2 MSC line and differentiated derivatives. Using undifferentiated C3H cells (U), together with differentiated adipocytes (A) and myocytes (M), we isolated three subcellular fractions for proteomic analysis: NE, nuclear contents (NC) and cytoplasmic membranes (CM) (Fig. 1; Materials and Methods). The NE and NC fractions were obtained by nuclease digestion of isolated nuclei followed by treatment with 0.5 M NaCl and sedimentation to yield the NE (pellet) and NC (supernatant) fractions. The CM fraction was obtained by flotation of membranes from a postnuclear supernatant to a low-density zone of a sucrose gradient, a procedure that enriches for secretory pathway organelles (Golgi, plasma membrane, and endosomes/lysosomes). We optimized our cell lysis and fractionation methods using Western blotting to follow marker proteins for various organelles (see Materials and Methods). The proteomics analysis of the fractions (below) provided a detailed measure of the relative abundance of benchmark and contaminant proteins in each fraction.

**Figure 1.**
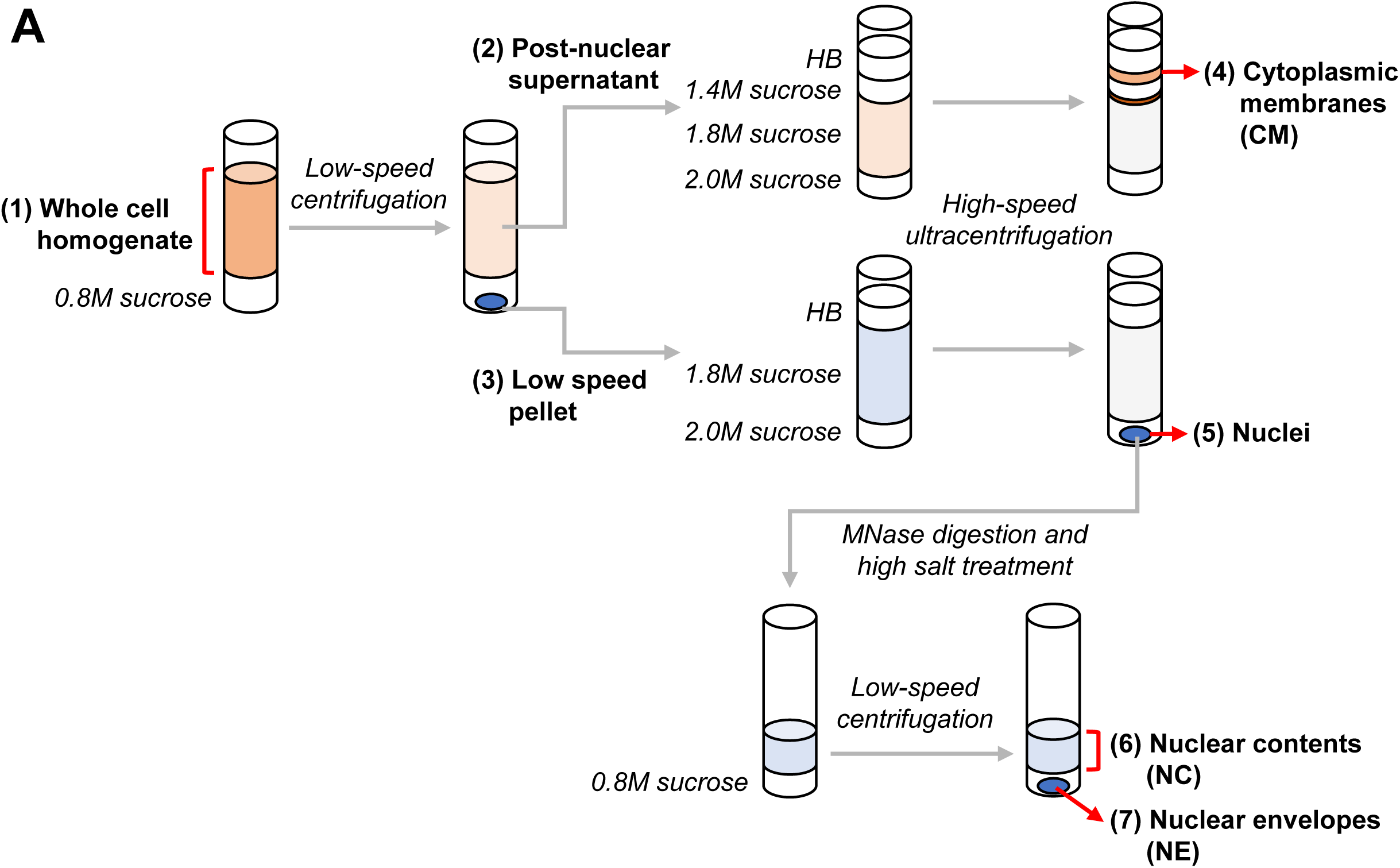
Isolation of subcellular fractions from U, A and M cells. Schematic diagram of the subcellular fractionation methods used to isolate NE, NC and CM fractions.

We used multidimensional protein identification technology (MudPIT [20]; see Materials and Methods) for proteomic analysis of the fractions from the three cell types. Collectively our analysis involved 3-4 mass spectrometry runs for each fraction and cell type, and identified 7938 proteins (Table S1). Approximately 60% of these were detected in all three cell types, whereas ~6-8% were uniquely found in only one of the three cells (Fig. 2A; Table S1). As expected, proteins diagnostic of differentiated adipocytes (e.g. long chain fatty acid CoA ligase 1, perilipin 1) and myocytes (myosin 3 heavy chain, titin) were strongly induced in the respective differentiated cells, based on NSAF (normalized spectral abundance factor [21]) values (Table S1). To evaluate the abundance of individual proteins in the NE fraction relative to NC and CM, we calculated a NE enrichment score (termed “score” below) based on NSAF values (see Materials and Methods). With this method, proteins that were detected only in the NE fraction had a score of 1, proteins that were found only in NC and/or CM had a score of 0, and proteins found both in the NE and in other fractions had intermediate scores. Scores were calculated only for proteins that were detected with 5 or more spectral counts in a particular cell type, to minimize less reliable predictions.

**Figure 2.**
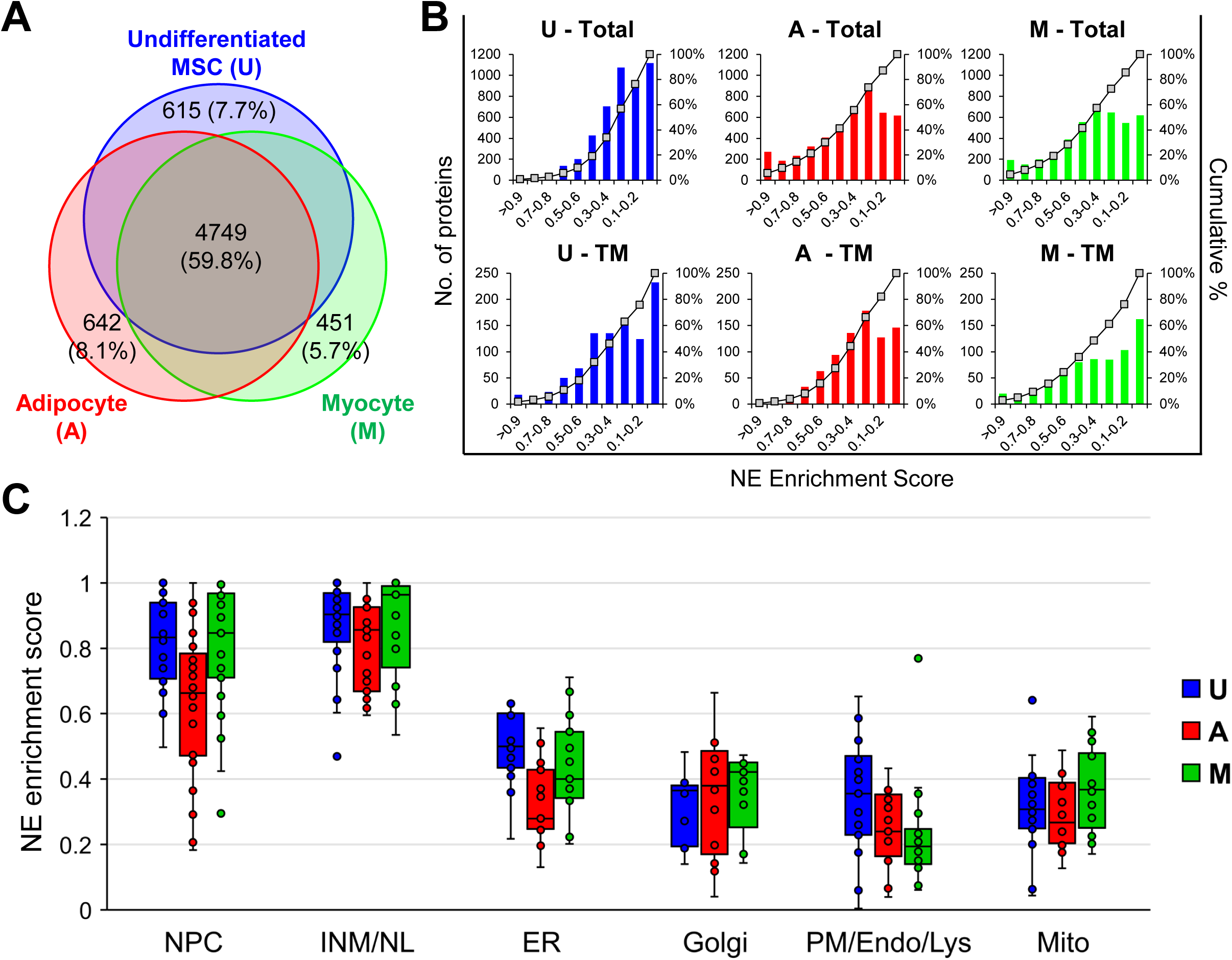
Analysis of overall results from the proteomic analysis. (A) Venn diagram representing proteins detected in U, A and M cells and the overlap of the datasets. (B) Graphs plotting the NE enrichment scores vs number of proteins in U, A and M cells for datasets representing all proteins (Total) or datasets representing only annotated TM proteins (TM). (C) Box and whisker plots depicting NE enrichment scores for benchmark proteins of the NE, NL/INM and the cytoplasmic membrane compartments indicated in U, A and M cells. See Table S2 for scores associated with specific proteins.

Only 3% of the proteins in U cells had scores ≥ 0.7 (Fig. 2B, top). By contrast, ~15-20% of the proteins scored in this range in the A and M cells. When only proteins with annotated TM segments were considered (roughly 20% of all proteins detected), the protein percentage scoring ≥ 0.7 was more similar in the three cell types (between 4-9%; Fig. 2B, bottom). Thus, the high-scoring protein set of U cells is relatively enriched in TM proteins, as compared to those of A and M cells. These differences are correlated with higher levels of tubulin in A and M cells and less efficient extraction of non-TM cytoplasmic and intranuclear proteins from the NE fraction of these cells, as compared to U.

We evaluated our scoring system using sets of benchmark proteins with well-defined membrane localizations (Table S2). The NE benchmarks, comprising 30 Nups and 23 INM proteins, included 22 proteins with TM segments. Other benchmarks involved sets of ~15-20 abundant TM proteins enriched in different cytoplasmic membrane compartments: peripheral ER, Golgi, mitochondria and the plasma membrane/endosome/lysosome system (Fig. 2B and Table S2). Most NE proteins with TM segments had a high score (> 0.7) in all three cell types. Conspicuous exceptions with lower scores (~0.5-0.7) were emerin and Tmem43 (LUMA), which are known to be partially localized to the peripheral ER in certain cell types and/or physiological states [12,13,22,23]. The benchmark TM proteins of the peripheral ER had scores ranging from 0.2-0.7, clustering around a mean value of ~0.5, although a few proteins characteristic of sheet ER (e.g. Sec11α, Sec61β) [24], had higher scores (~0.7-0.8) in one or more cell types. TM proteins of Golgi, plasma membrane/ endosome/lysosome and mitochondria mostly had scores between 0.1-0.5, consistent with Western blot analysis of the NE fractionation (data not shown).

Many of the benchmark NE proteins lacking annotated TM segments also had high scores (> 0.7) in the three cell types, including B-type lamins, the NL-associated proteins Prr14 [25] and Gmcl1 [26], many Nups and the NPC-associated protein Mcm3ap [27]. NE proteins with somewhat lower scores included A-type lamins and ~10 Nups lacking TM segments, which correspondingly were relatively abundant in the NC fraction. The latter results are consistent with the well-established existence of intranuclear pools of lamins A/C [28] and Nups [29] separate from the NL and NPC, respectively. The scores for almost all NE markers in A were lower than their corresponding scores in U and M cells (Table S2), coinciding with relatively higher levels of these proteins in the NC fraction (Table S1). This may reflect greater fragility of the NE of A cells, resulting in release of NE fragments to NC during fractionation.

The non-TM protein datasets included low-abundance components with high scores in one or two cell types. Many of these have known regulatory, enzymatic or structural roles in the nuclear interior, cytosol or extracellular matrix. Accordingly, we consider it likely that most proteins in this group are not concentrated at the NE *in situ*. The high scores could reflect either intrinsic limitations of MudPIT proteomics (see Discussion), adsorption to the NE during isolation, or association with co-fractionating structures such as the intermediate filament protein synemin or the extracellular matrix components collagen or fibronectin.

The above considerations indicate that our scoring system can most effectively predict new NE-concentrated protein with TM segments. We performed unsupervised hierarchical clustering to further analyze the set of 243 TM proteins with scores higher than 0.5, sorting the proteins from the three cell types into eight clusters (Fig. 3A and Table S3). The cluster with high scores in all three cell types (Fig. 3A) was predominated by well-established NE-concentrated proteins (see Table S2). It also included another five proteins not known to be concentrated at the NE, which represent new candidates. We selected four members of this group to analyze: Arl6ip6, Mfsd10, Smpd4 and Tmx4 (Fig. 3B). We added a fifth high-scoring protein found in the three cell types (Itprip) to the query group, since it is predicted to have a TM domain by the CCTOP consensus algorithm and is homologous to Itpripl1 and Itpripl2, which both contain a curated TM segment. We also analyzed another member of the cluster that recently was shown to be concentrated at the NE, Vrk2 [30]. A diagrammatic representation of these proteins, with the position of the epitope tag and predicted TM segments, is shown in Fig. 3C. The specific peptides detected for these proteins are listed in Table S3.

**Figure 3.**
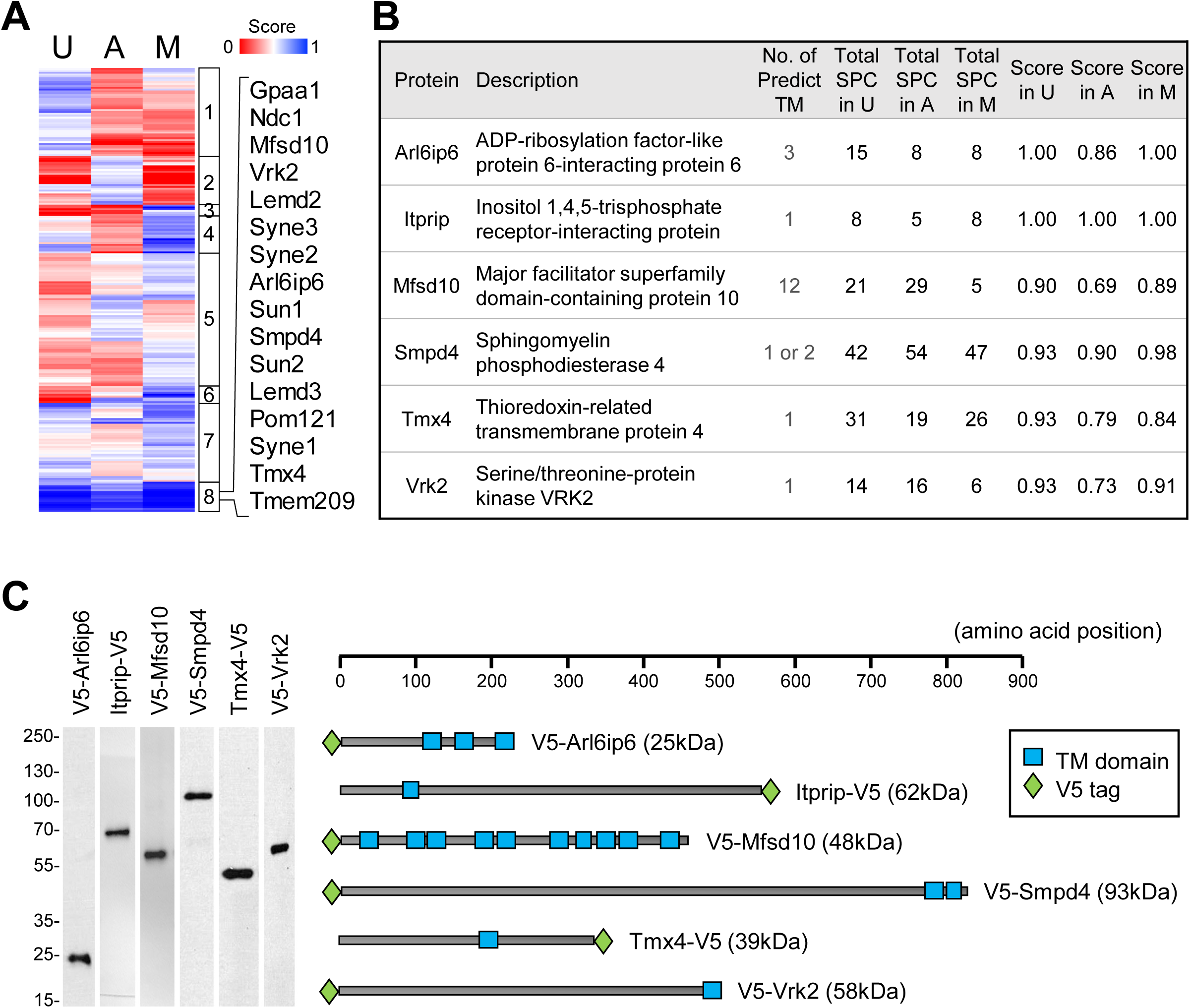
Cluster analysis and selection of target set. (A) Depiction of cluster analysis of annotated TM proteins in the three cell types. Right panel: List of proteins in cluster 8. (B) Proteins selected for analysis, together with a description of each protein, the number of annotated TM domains, spectral counts and NE enrichment score obtained in each cell type. (C) Left: Western blot analysis of C3H cell populations stably expressing each of the candidates. Right: Depiction of the size of the proteins, the position of the epitope tag, and predicted transmembrane segment(s).

To evaluate whether these candidates are concentrated at the NE relative to the peripheral ER, we prepared populations of C3H cells transduced with lentiviral vectors expressing epitope-tagged versions of the proteins. We also generated control cell strains expressing the NE proteins Lem2 and emerin, and the pan-ER marker Sec61β [24]. After verifying that the ectopic proteins migrated at their predicted sizes on SDS gels (Fig. 3C), we carried out immunofluorescence microscopy to detect the tagged proteins and two endogenous markers, lamin A and the pan-ER TM protein calnexin (Fig. 4 and Figs. S1-S2). We found that all five of our candidates were substantially concentrated at the NE relative to calnexin and Sec61β, similar to ectopically expressed Lem2 and emerin. The ectopic constructs also were detected in cytoplasmic areas at relatively low levels that varied from cell to cell (Fig. 4 and Fig. S2). Smpd4 appeared in cytoplasmic foci of varying sizes (Fig. 4 and Fig. S3D), whereas the remaining four candidates closely co-localized with peripheral ER calnexin. Vrk2 was clearly concentrated at the NE relative to calnexin, but it appeared in the peripheral ER at substantially higher levels than the other ectopic constructs. Previous work also revealed substantial peripheral ER labeling of ectopic Vrk2 in stably expressing cells [30].

**Figure 4.**
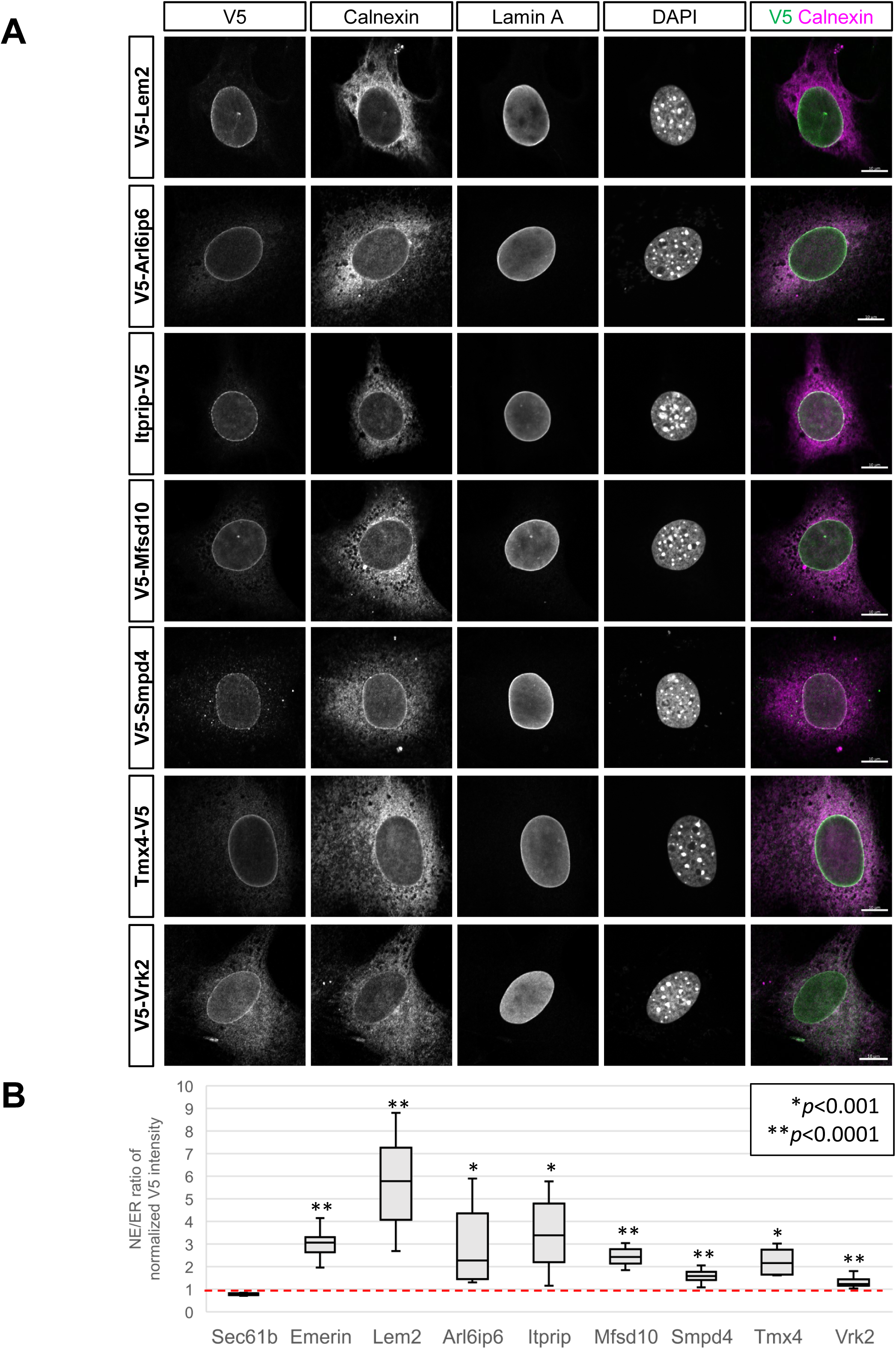
Immunofluorescence localization of ectopically expressed target proteins. (A) Micrographs of representative cells from ectopically expressing populations that were co-labeled with antibodies to the V5 epitope tag (to detect indicated targets), calnexin and lamins A/C. DNA staining (DAPI) and a merge of V5 and calnexin labeling is shown in the right panels. Lem2 and Vrk2 are included as positive controls. Scale bars, 10μm. (B) Graph showing relative enrichment of targets in the NE, determined by ratio of the level of V5/calnexin staining in the NE vs the V5/calnexin ratio in the immediately adjacent peripheral ER. Quantification is described in Materials and Methods. Red line indicates the V5/calnexin ratio in the NE vs peripheral ER for ectopically expressed Sec61β. Stably transduced populations were examined in all cases except for Smpd4, which was visualized in transiently transduced cells (see Materials and Methods).

During the course of this analysis, it became evident that expression levels influenced the relative concentration of ectopic proteins in cytoplasmic vs NE regions, particularly for Tmx4 and Smpd4. Whereas Tmx4 was highly concentrated at the NE compared to the peripheral ER in populations expressing lower protein levels, no NE concentration was evident in cells expressing higher levels (Fig. S3A-C). Also, the selective NE targeting of ectopic Smpd4 that was evident with low expression at early times after lentiviral transduction (Fig. 4 and Fig. S3D was largely obscured by numerous cytoplasmic Smpd4 foci that accumulated in stably expressing cells (Fig. S3D). These results are consistent with the diffusion-retention model for NE localization, since saturation of NE binding sites by ectopic overexpression is expected to result in protein redistribution to the contiguous peripheral ER or its appearance in aggregates. Accordingly, we analyzed cells reflecting the lower half of the expression spectrum for the determination of NE enrichment in Fig. 4.

We implemented an unbiased quantitative method to determine the relative concentration of the ectopic constructs at the NE. This involved measuring the fluorescence intensity ratio for the epitope tag/endogenous calnexin at the NE and adjacent peripheral ER around the circumference of the nucleus in cross-section. We defined the NE by the concentrated nuclear rim of lamin A staining, and the peripheral ER by a concentric zone located 0.5-1 μm outside the NE. Using this method, we determined that ectopically expressed emerin and Lem2 were ~3-6-fold concentrated at the NE. The five NE candidates were 1.7-3.5-fold concentrated at the NE, whereas Vrk2 showed a lower (1.3-fold) but statistically significant NE concentration. For the reasons considered in the above paragraph, the peripheral ER levels seen with ectopically expressed constructs might over-represent the native levels of these proteins in the peripheral ER. Unfortunately, we were unable to determine the ectopic/endogenous expression levels for any of the constructs (see Materials and Methods).

We used cell permeabilization with low/high concentrations of digitonin to analyze whether the new NE-concentrated proteins are exposed to the ONM, or reside in a sequestered space at the INM or NPC (Fig. 5A). With this technique, low concentrations of digitonin permeabilize the plasma membrane and allow antibody access to the cytosolic space and ONM, but leave the ER and NE intact [31]. Conversely, high concentrations of digitonin fully permeabilize the NE and allow antibody access to proteins of the INM and NPC-associated membrane as well. We validated this method in C3H cells by antibody labeling of calnexin and Lem2 (Fig. S5): only the cytosolically exposed epitopes recognized by the calnexin antibody were accessible with low digitonin, but both calnexin and ectopically expressed Lem2 (concentrated at the INM) were labeled after cell treatment with either high digitonin or Triton X-100. When applied to analysis of the five new NE proteins and Vrk2, low digitonin treatment yielded strong NE staining only for Itprip, although labeling of the low-level peripheral ER pools of Arl6ip6, Mfsd10, Tmx4 and Vrk2 also was evident. Treatment with high digitonin yielded strong NE labeling of the latter four proteins, as well as strong staining of Smpd4 at both the NE and in cytoplasmic foci. These results indicate that Itprip is exposed to the ONM, and suggest that the remaining proteins are located at membrane-sequestered NE regions. Since Arl6ip6, Mfsd10 and Tmx4 showed relatively uniform nuclear rim staining similar to Vrk2, it is likely that these proteins are localized at the INM.

**Figure 5.**
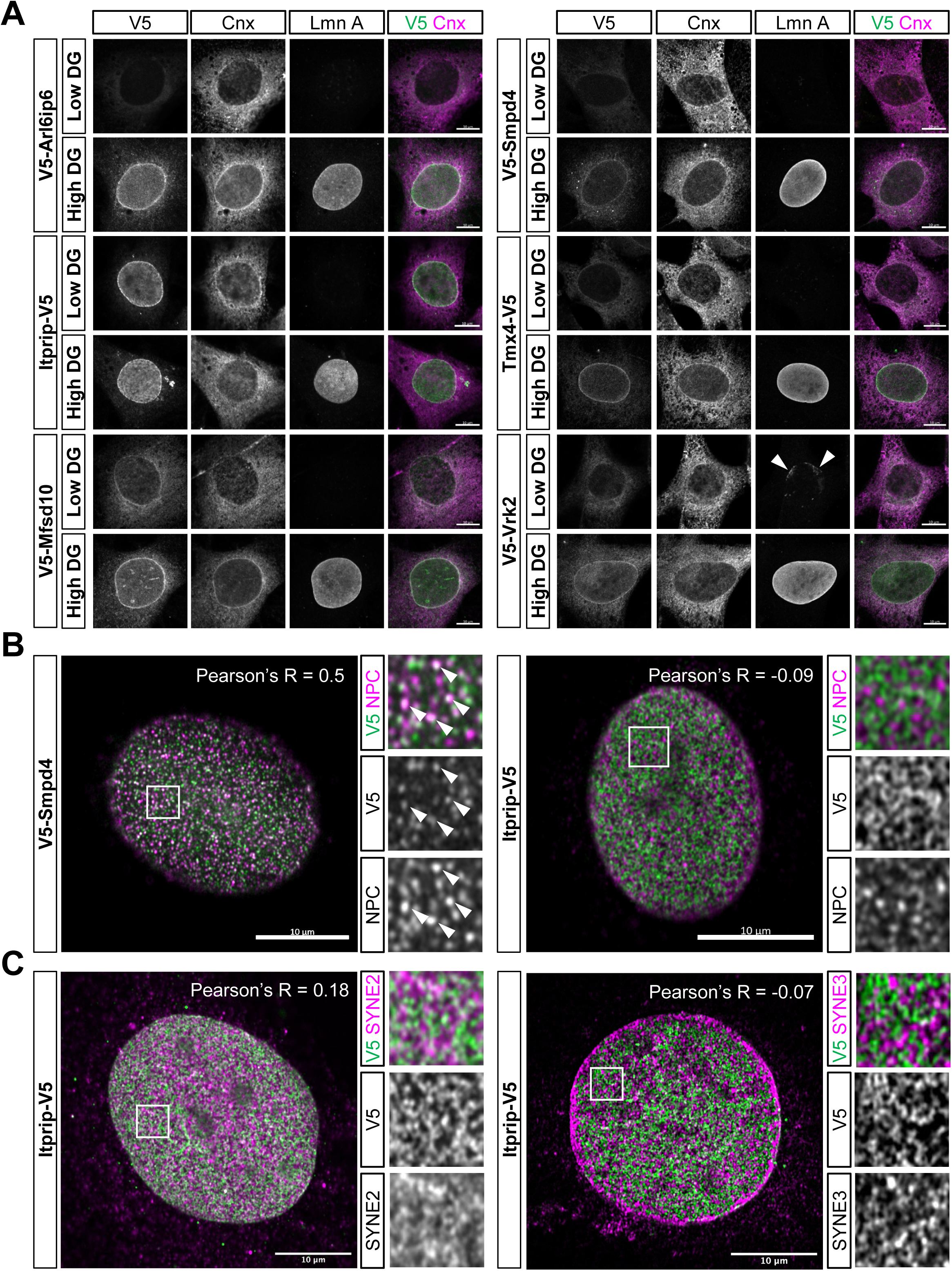
Immunofluorescence analysis of the localization of target proteins with respect to NE substructure. (A) Representative images obtained with antibody staining after cell permeabilization with low (low DG) or high (high DG) concentrations of digitonin. C3H cells expressing the depicted ectopic targets were incubated with antibodies to the V5 tag, calnexin and lamin A. Merged image of V5 and calnexin staining is shown on the right. (B, C) High resolution immunofluorescence images of cells expressing ectopic Smpd4 or Itprip as indicated. Images were obtained with Airyscan (Materials and Methods) using cells fixed and permeabilized with standard conditions. (B) Cells were co-labeled with antibodies to V5 and RL1, a monoclonal antibody recognizing FG repeat Nups of the NPC [63]. Pearson’s correlation coefficient R (indicated) shows moderate co-localization between V5-Smpd4 and the NPC but no significant co-localization between Itprip-V5 and the NPC. Right panels show higher magnification views of areas indicated by boxes. In cells expressing V5-Smpd4, examples of foci co-labeled with RL1 and anti-V5 are indicated (arrowheads). (C) Cells were co-labeled with antibodies to V5 and either nesprin-2 (Syne2) or nesprin-3 (Syne3). Right panels show higher magnification views of areas indicated by boxes. Pearson’s correlation coefficient R (indicated) shows no significant co-localization between Itprip-V5 and nesprin-3, and weak co-localization between Itprip and nesprin-2. All panels: scale bars, 10μm. Stably transduced populations were examined in all cases except for Smpd4, which was visualized in transiently transduced cells.

The NE staining seen for both Smpd4 and Itprip was conspicuously less uniform than that of the other proteins analyzed. Using high resolution imaging to capture surface views of the NE (Materials and Methods), we found that Smpd4 was localized to small puncta at the nuclear surface (Fig. 5B and Fig. S4). In substantial part, these puncta co-localized with NPCs, as detected by an antibody to FG-repeat Nups (Pearson correlation coefficient R = 0.5). However, the Smpd4 intensity in different NPC puncta varied considerably, and some of the Smpd4 puncta at the NE had little or no Nup staining. Surprisingly, many of the cytoplasmic Smpd4 foci seen in transiently transduced cells, which were present with much higher abundance in stably expressing populations, also were strongly labeled with the antibody to FG Nups (Fig. S3D. This suggests that FG Nups are recruited to ectopic cytoplasmic foci containing Smpd4. A similar pattern was reported with ectopic overexpression of the transmembrane Nup Pom121, which induced cytoplasmic foci containing both Pom121 and FG Nups [32]. These results, together with data revealing interactions of Smpd4 with several Nups in pull-down assays [33], support a physiologically relevant interaction between Smpd4 and Nups, and strongly suggest that at least much of Smpd4 is associated with the NPC. The non-uniform co-localization of Smpd4 and FG Nups at the nuclear surface in part could reflect uneven association of ectopic Smpd4 with different populations of NPCs. It also may be a consequence of Smpd4 overexpression and assembly into non-native NPC-related structures at the nuclear surface.

The distribution of Itprip on the nuclear surface was qualitatively different from the Smpd4 staining (Fig. 5B-5C and Fig. S4), appearing as linear arrays of puncta instead of the more distributed NPC-like pattern. The Itprip puncta did not co-localize with FG Nups (Fig. 5B, nesprin-1 (data not shown), or nesprin-3 (Fig. 5C). We did observe weak co-localization with nesprin-2 (Fig. 5C), although more analysis will be required before conclusions can be made on this potential association.

We next analyzed whether the five newly identified NE proteins and Vrk2 are concentrated at the NE in differentiated adipocytes and myocytes (Fig. 6), as suggested by their high proteomics scores. We were able to visualize Arl6ip6, Itprip, Smpd4, Tmx4 and Vrk2 in differentiated adipocytes and in all cases obtained strong labeling of the NE with little or no peripheral ER staining. We achieved myogenic differentiation of cells stably transduced with Tmx4, Itprip and Arl6ip6, but not with the other proteins (see Materials and Methods). In all three cases, we observed robust NE-concentrated staining. As an additional model for NE targeting, we analyzed stably transduced populations of the human U2OS osteosarcoma cell line. In all cases, we observed strong targeting of the new proteins to the NE (Fig. S6). Together these results indicate that the newly characterized proteins have the capacity to concentrate at the NE in a variety of different cell types, and likely are widespread NE components.

**Figure 6.**
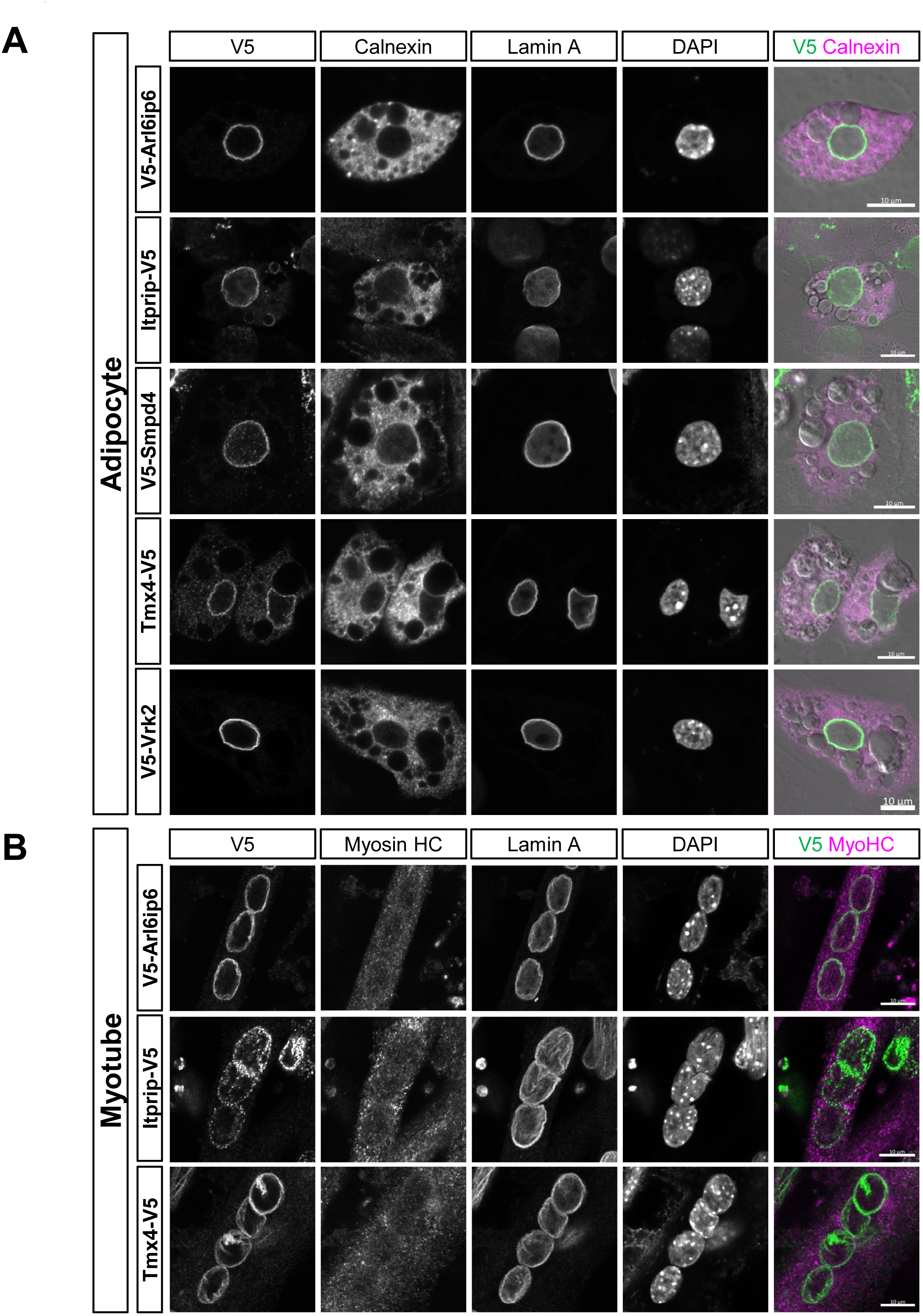
Immunofluorescence analysis of the localization of target proteins in adipocytes and myocytes. (A) C3H cells stably transduced with the indicated constructs were differentiated into adipocytes and co-labeled with antibodies to the V5 epitope tag, calnexin and lamins A/C. DNA staining (DAPI) and a merge of V5 and calnexin labeling is shown in the right panels. (B) C2C12 myoblasts that were stably transduced with the indicated constructs were differentiated into myotubes and labeled as in (A). Scale bars, 10μm.

## Discussion

Here we used MudPIT analysis of subcellular fractions to identify five previously unrecognized TM proteins that are strongly concentrated at the NE: Arl6ip6, Itprip, Mfsd10, Smpd4 and Tmx4. Immunofluorescence microscopy with epitope accessibility analysis revealed that Itprip is located at the ONM and that much of Smpd4 is concentrated at the NPC. The experiments also suggested that the other three proteins reside in the INM (Fig. S7). Although we focused our analysis on mesenchymal stem cells, adipocytes and myocytes, we found that these proteins also target strongly to the NE in U2OS cells. RNA-seq databases indicate that the five proteins are expressed broadly in human and mouse tissues, and all have been detected in HeLa cells by proteomics [33]. Thus, these probably are widespread NE components.

The proteomics scoring system we used to evaluate NE enrichment was validated by examination of benchmark proteins for several membrane compartments, and provided a strong framework for our efforts. One caveat of this strategy is that MudPIT proteomics only semi-quantitatively represents the relative abundance of a particular protein in different subcellular fractions, with higher spectral detection increasing the reliability [34]. Thus, scores based on relatively low spectral counts should be interpreted cautiously.

We found that validation of NE proteins by ectopic expression was most accurately achieved using stably transduced cell populations, except in the case of Smpd4 (see Materials and Methods). With these approaches, we deem it essential to quantify the relative concentration of the ectopic target at the NE vs the peripheral ER using an internal TM marker that is evenly represented in both membrane systems, such as calnexin. Our quantification method involved Image J software-based analysis of protein concentration around the nuclear circumference using lamin A to define the position of the NE. We emphasize that high levels of ectopic protein overexpression in some cases can mask NE localization, as we have observed for Tmx4. This can explain the discrepancy between our demonstration that Tmx4 is concentrated at the NE with relatively low ectopic expression, and previously published work revealing a pan-ER distribution of ectopic Tmx4 [35,36], similar to our results with high Tmx4 expression.

Aside from the sample set we analyzed, we consider it likely that some additional TM proteins with high scores in our analysis will turn out to be NE-concentrated, even in cases where database annotations suggest otherwise. For example, Pigb (not detected in our analysis) is known to be a mannosyltransferase involved in synthesis of the GPI anchor precursor and is annotated in UniProtKB as a general ER protein, but was recently shown to be strongly concentrated at the NE in *Drosophila* [37]. Here we analyzed proteins with high NE enrichment scores in all three cell types, which appeared in one of the clusters. Some of the proteins in other clusters, which have high scores in one or two of the cell types, might be concentrated at the NE is a differentiation state-selective pattern.

The group of high-scoring non-TM proteins in our datasets also is likely to contain proteins concentrated at the NE in mesenchymal cells. NE association has been suggested for some of these, such as Akap8l [38], the prostaglandin synthase Ptgs2 [39] and the choline phosphate cytidylyltransferase Pcyt1a [40,41]. However, we consider our datasets and scoring system most useful for the analysis of TM proteins.

Sequence analysis of the newly identified NE proteins suggests potential roles in nuclear regulation and membrane dynamics. The most evolutionarily conserved protein of this group is Mfsd10, a member of the ancient Major Facilitator Superfamily of membrane solute transporters. Mfsd10 was proposed to be a cellular efflux pump for organic anions and nonsteroidal anti-inflammatory drugs [42], although its transport properties have not been directly analyzed. Interestingly, the highest Psi-BLAST scores for Mfsd10 involve tetracycline efflux pumps of gram negative bacteria (e.g. 31% identity/45% similarity over 90% of Mfsd10 sequence with the TetA(41) gene of S. *marcescens)*. This raises the possibility that Mfsd10 may transport toxic metabolites and/or xenobiotics across the INM to the ER lumen, as a means of efficiently funneling deleterious compounds out of the nuclear environment.

Unexpectedly, we found the sphingomyelin phosphodiesterase Smpd4 [43] to be concentrated at the NPC. Smpd4 releases ceramide, a signaling molecule itself and biosynthetic precursor to the signaling lipid S1P [44] that is known to act in the nucleus to inhibit HDACs [45,46] and to stabilize telomerase [47]. The NPC association of Smpd4 raises the possibility that the production of S1P in the nucleus might be linked to transport activity at the NPC. Smpd4 also might have a role in lipid bilayer dynamics at the NE. For example, if Smpd4 were localized on the INM side of the NPC, sphingomyelin hydrolysis could reduce lipid packing on the nucleoplasmic leaflet of the INM to drive the membrane association and activation of Pcyt1a [40] or the pro-inflammatory phospholipase A2 [48]. Sphingomyelin hydrolysis also could promote local concave membrane curvature, which accompanies the process of NPC insertion in the interphase NE [49].

Itprip was the only protein of the group found to be localized to the ONM. Its concentration at the NE could be explained most simply by an interaction with one or more nesprins, which themselves are concentrated at the NE due to transmembrane associations with SUN-domain proteins [15,16]. We detected weak co-localization of Itprip with nesprin-2 and no co-localization with nesprin-1 or nesprin-3 using the antibodies that were available, but further analysis is needed to investigate these potential associations. Previous studies have shown that Itprip binds the inositol triphosphate receptor calcium channels and negatively regulates their activity *in vitro* [50]. Thus, regulation of calcium fluxes at the NE is a potential function of Itprip at the NE. In one scenario, this could be connected to mechanical communication between the cytoskeleton and the LINC complex.

The properties of Tmx4 and Arl6ip6 are consistent with a role in regulating NE structure. The thioredoxin domain of Tmx4 is likely localized to the NE lumen, since it occurs between an N-terminal signal sequence and the single TM segment. This suggests a potential role in regulating the luminal aspects of NE specific structures, such as the LINC complex and associated torsinA, both of which may be regulated by disulfide oxidation/reduction [51,52]. Arl6ip6 is a susceptibility locus for ischemic stroke [53]. It lacks enzyme-related domains, but does show physical interactions in proteome-wide pull-down screens with a number of proteins involved in membrane vesicle formation/targeting [33,54], a process that is involved in NE resealing and repair.

In conclusion, the set of new NE proteins identified in this study provide new avenues for studying the dynamics and functions of the NE. It will be useful to extend the methodology used in this study to the analysis of other cell types, where we expect that additional, interesting NE-concentrated proteins will be identified.

## Materials and Methods

### Cell culture

C3H/10T1/2 (C3H), C2C12, U2OS, and 293T cells were all acquired from the American Type Culture Collection (ATCC). C3H, C2C12, and 293T cells were grown in high glucose Dulbecco’s modified Eagle’s medium (DMEM) (Gibco) supplemented with 10% fetal bovine serum (FBS) (Gibco), 1% penicillin/streptomycin/glutamine cocktail (P/S/G) (Gibco), and 1% minimum essential medium non-essential amino acids (NEAA) (Gibco). U2OS cells were grown in McCoy’s 5A medium (Gibco) supplemented with 10% FBS and 1% P/S/G. All cells were maintained at 37°C in 5% CO2.

### Cell Differentiation

For adipogenesis, C3H cells were grown on plates coated with 0.2% gelatin (Sigma) and allowed to reach 100% confluency. Growth medium was then changed to C3H differentiation medium consisting of DMEM supplemented with 10% iron-supplemented bovine calf serum (HyClone) and 1% P/S/G. After 48 hours, medium was changed to pre-adipocyte differentiation medium consisting of DMEM-L-Glutamax (Gibco) supplemented with 10% FBS (Gemini), 1% P/S/G, 5 μg/mL human insulin (Sigma), 0.5 mM 3-isobutyl-1-methylxanthine (IBMX) (Sigma), 1 μM Dexamethasone (Sigma), and 2 μM Rosiglitazone (Cayman). After another 48 hours, media was changed to adipocyte differentiation medium consisting of DMEM-L-Glutamax supplemented with 10% FBS (Gemini), 1% P/S/G, and 5 μg/mL human insulin. Adipocyte differentiation medium was replaced every 2 to 3 days until terminal differentiation (5 to 7 more days).

For C3H myogenesis, cells were grown in DMEM supplemented with 10% FBS and 1% P/S/G. C3H cells were transfected with 10 μg of a doxycycline-inducible MyoD piggyBac transposon vector [55]. After 48 hours, cells with positive integration of the vector were selected using 2 μg/mL puromycin (Invivogen) for 24 hours. To initiate myotube differentiation, stably integrated populations of C3H cells were grown on 500 cm plates to 70% confluency and induced with 20 ng/mL doxycycline (Sigma) for 24 hours. Cells were then changed to differentiation medium containing DMEM with 2% donor equine serum (HyClone), 1% IPS solution (Sigma), and 1% P/S/G. Differentiation media was replaced every 24-48 hours until terminal differentiation (about 3 days).

For C2C12 myogenesis, cells were plated and allowed to reach 90-95% confluency. Media was then changed to DMEM supplemented with 1% donor equine serum (HyClone), 1% P/S/G, and 1% ITS Liquid Media Supplement (Sigma). Medium was replaced every 48 hours until terminal differentiation (4 to 5 days after initiation of myogenesis).

### Subcellular fractionation

For subcellular fractionation, C3H cells were seeded in 500 cm^2^ plates and allowed to reach 90% confluency. Plates were rinsed three times with ice-cold PBS, and then three times with ice-cold homogenization buffer (HB) (10 mM HEPES pH 7.8, 10 mM KCl, 1.5 mM MgCl_2_, 0.1 mM EGTA) containing 1 mM DTT, 1 mM PMSF, and 1 μg/mL each of pepstatin, leupeptin, and chymostatin. After these washes, cells were incubated in HB for 15 minutes on ice. Cells were then scraped off plates and were further disrupted by dounce homogenization with 18-20 strokes. The whole cell homogenate was then layered on top of 2 mL shelf of 0.8 M sucrose in HB and centrifuged at 2000 rpm for 10 minutes at 4°C in a JS5.2 rotor with no brake to yield a crude nuclear pellet and postnuclear supernatant. The postnuclear supernatant comprising the zone above the sucrose shelf, and pelleted nuclei were each resuspended in 1.8 M sucrose (final concentration) in HB using a cannulus. The resuspended nuclei and postnuclear supernatant were layered in separate ultra-clear 13.2 ml nitrocellulose centrifuge tubes on top of a 1 mL layer of 2.0 M sucrose in HB. For the nuclear gradient, HB was layered over the loading zone to fill the nitrocellulose tube. For the postnuclear supernatant gradient, 1 mL of 1.4 M sucrose in HB was layered on top of the loading zone, followed by HB to fill the tube. The gradients then were centrifuged at 35,000 rpm (210,000g) for 1 hour at 4°C with no brake in an SW41Ti rotor. Nuclei that pelleted through the 2.0 M sucrose were resuspended in HB and dounce homogenized with 2 strokes to disperse aggregates. For the postnuclear supernatant gradient, the HB/1.4 M sucrose interphase was collected and saved as “cytoplasmic membranes” (CM). Nuclei were then incubated with 1 mM CaCl_2_ and 100 ku/mL micrococcal nuclease (New England Biolabs) in HB for 37°C for 15 minutes. Digested nuclei were then placed on ice and NaCl was added to a final concentration of 500 mM. The digested nuclei sample was layered on top of 1 mL shelf of 0.8 M sucrose in HB and centrifuged at 4000 rpm for 10 minutes at 4°C in a JS5.2 rotor. A sample comprising the region above the 0.8 M sucrose layer was collected and saved as “nuclear contents” (NC). The NE fraction, comprising the pellet, was collected by resuspension in HB. During development of the fractionation method, we monitored different organelles and cellular components at progressive steps of the isolation with antibodies to the following markers: lamin B1 and the INM resident LAP2β for the NE, histone H2B for chromatin, calnexin for sheet and tubular ER [24], Tim23 for mitochondria and Pex14 for peroxisomes.

### MudPIT Proteomics

30 μg protein from each subcellular fraction (NE, NC and CM), estimated by the Pierce BCA protein assay (Thermo Fisher), was made up to a final concentration of 4 M urea, 0.2% RapiGest SF (Waters Corporation) and 100 mM NH_4_HCO_3_ pH 8.0. Proteins were reduced with Tris(2-carboxyethyl)phosphine hydrochloride and alkylated with 2-Chloroacetamide. Next, proteins were digested with 0.5 μg Lys-C (Wako) for 4 hours at 37°C, and then for 12 hours at 37°C in 2 M urea, 0.2% RapiGest SF, 100 mM NH_4_HCO_3_ pH 8.0, 1 mM CaCl_2_ with 1 μg trypsin (Promega). Digested proteins were acidified with TFA to pH < 2 and RapiGest SF was precipitated out. Each fraction was loaded on individual MudPIT micro-columns (2.5 cm SCX: 5 μm diameter, 125 Å pores; and 2.5 cm C18 Aqua: 5 μm diameter, 125 Å pores; Phenomenex), and resolved across an analytical column (15 cm C18 Aqua: 5 μm diameter, 125 Å pores) (Phenomenex).

Analysis was performed using an Agilent 1200 HPLC pump and a Thermo LTQ-Orbitrap Velos Pro using an in-house built electrospray stage. MudPIT experiments were performed with steps of 0%, 10%, 20%, 30%, 50%, 70%, 80%, 90%, 100% buffer C and 90/10 % buffer C/B [20], being run for 5 min at the beginning of each gradient of buffer B. Electrospray was performed directly from the analytical column by applying the ESI voltage at a tee (150 mm ID) (Upchurch Scientific) [20]. Electrospray directly from the LC column was done at 2.5 kV with an inlet capillary temperature of 325^o^C. Data-dependent acquisition of tandem mass spectra were performed with the following settings: MS/MS on the 20 most intense ions per precursor scan; 1 microscan; reject unassigned charge state and charge state 1; dynamic exclusion repeat count, 1; repeat duration, 30 second; exclusion list size 500; and exclusion duration, 90 second.

Protein and peptide identification was done with the Integrated Proteomics Pipeline - IP2 (Integrated Proteomics Applications, Inc. http://www.integratedproteomics.com/). Tandem mass spectra were extracted (monoisotopic peaks) from raw files using RawConverter [56] and were searched against a UniProt SwissProt *Mus musculus* database (release 2014_01) with reversed sequences using ProLuCID [57,58]. The search space included all fully-tryptic and half-tryptic peptide candidates with static modification of 57.02146 on cysteines. Peptide candidates were filtered using DTASelect [59] at 1% protein level False Discovery Rate, (parameters: -p 1 -y 1 – trypstat –pfp 0.01 –extra –pI -DM 10 –DB –dm -in -t 0 –brief –quiet) [60,61].

To calculate a NE enrichment score, the sums of the NSAF scores for each protein for NE, CM, and NC fractions were calculated. The following equation was used to determine NE enrichment score, where e = experimental run and p = protein ID:

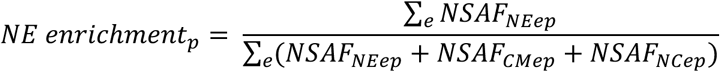

The clustering analysis was carried out only on annotated TM proteins with an enrichment score of greater than 0.5 in at least one of the three cell types (U, A, M). These proteins then were clustered on the basis of their enrichment scores in all three cell types. An unsupervised hierarchical clustering algorithm was applied using the Euclidean distances of the score triplets using the *“hclust’* R function with the *“complete”* agglomeration method. 8 clusters were selected and plotted in Figure 3A and Table S3.

The proteomics datasets have been deposited in the public proteomics repository MassIVE (Mass Spectrometry Interactive Virtual Environment), part of the ProteomeXchange consortium [62], with the identifier MSV000083166 (and PXD011856 for ProteomeXchange) and is available through the following link: ftp://massive.ucsd.edu/MSV000083166.

### Antibodies

The following primary antibodies were used for Western blotting: Rabbit anti-Calnexin (Sigma #C4731), Rabbit anti-Lamin B1 (made in-house), Rabbit anti-LAP2β (made in-house), Rabbit anti-H2B(V119) (Cell Signaling #8135S), Mouse anti-Tim23 (BD Transduction Laboratories #611222), Rabbit anti-Pex14 (Millipore #ABC142). Unfortunately, we were unable to compare ectopic and endogenous levels of the five NE candidates, since none of the commercial antibodies we analyzed convincingly detected the endogenous counterparts by Western blotting of whole cell lysates.

The following primary antibodies were used for immunofluorescence staining: mouse anti-V5 (Invitrogen #46-0705), rabbit anti-V5 (Thermo Fisher #PA1-993), rabbit anti-calnexin (Abcam #ab22595), guinea pig anti-lamin A/C (made in-house), mouse anti-NPC RL1 (IgM, [63]), mouse anti-nesprin-1 (8C3, gift from Dr. Glenn E. Morris, RJAH Orthopaedic Hospital, UK), mouse anti-nesprin-1 (7A12, Millipore #MABT843), rabbit anti-nesprin-2 (Invitrogen OR United States Biological Corporation), rabbit anti-nesprin-3 (United States Biological Corporation), rabbit anti-myosin heavy chain (Abcam #124205). DAPI (Sigma) was used to stain DNA.

### Molecular cloning

To construct lentiviral vectors containing V5-tagged versions of our genes of interest, the following cDNA clones were purchased: pCMV6-Arl6ip6 (Origene, RefSeq BC019550), pCMV6-Tmx4 (Origene, RefSeq NM_029148), pCMV6-Mfsd10 (Origene, RefSeq NM_026660), pCMV6-Vrk2 (Origene, RefSeq NM_027260), pCMV6-Tmem128 (Origene, RefSeq NM_025480), pCMV6-Ccdc167 (Origene, RefSeq NM_001163741), pcDNA3.1-eGFP-Smpd4 (Genscript, RefSeq NM_029945), pcDNA3.1-ITPRIP-DYK (Genscript, RefSeq NM_001272012). mCherry-Sec61β was a gift from Gia Voeltz (Addgene plasmid #49155). Lem2 and Emerin were cloned from plasmids previously constructed in our lab.

All genes were inserted into pLV-EF1a-IRES-Puro (gift from Tobias Meyer, Addgene plasmid #85132) using ligation independent cloning (LIC). Two LIC-compatible sites, containing either an N-terminal V5 tag or a C-terminal V5 tag, were designed synthetically and inserted into pLV-EF1a-IRES-Puro using restriction enzymes BamHI and MluI. The primers used to insert the genes of interest into the LIC-compatible pLV-EF1a-IRES-Puro vector are listed in Supplementary Table S4. The portion of the primer that aligns to the gene of interest is underlined, and the portion that is required for T4 polymerase digestion during LIC is not underlined.

Genes of interest were amplified by PCR using Phusion High-Fidelity DNA Polymerase (New England Biolabs). pLV-EF1a-IRES-Puro LIC-compatible vectors were digested with SrfI (New England Biolabs). PCR fragments and SrfI-digested vector were treated with T4 Polymerase (New England Biolabs) in the presence of either dTTP (PCR products) or dATP (vector). T4P-digested vector and inserts were mixed at room temperature for 5 minutes, and then NEB Stable Competent Cells (New England Biolabs) were transformed with the product. Cells were incubated at 30°C for 24 hours, and then colonies were picked for clone validation. All cDNA clones were confirmed by complete DNA sequencing of the ORF in both 5’-3’ and 3’-5 directions.

### Generation of stably transduced cell populations

To produce lentiviruses, 293T cells were seeded so that they would be 60% confluent for transfection. Medium was changed to DMEM supplemented with 10% FBS, 1% glutamine, and 1% NEAA without antibiotics 30 minutes prior to transfection. Cells were transfected with pRSV-REV (gift from Didier Trono, Addgene plasmid #12253), pMDL-RRE (gift from Didier Trono, Addgene plasmid #12251), pCMV-VSVg (gift from Bob Weinberg, Addgene plasmid #8454), and pLV-EF1a-gene-of-interest (pLV-EF1a-GOI) vectors using Lipofectamine 2000 (Invitrogen). Viral supernatant was harvested 48 hours after transfection and filtered through a 0.45 μm polyethersulfone membrane filter (GE Healthcare Whatman).

C3H, C2C12, and U2OS cells were changed to DMEM with 10% FBS, 1% glutamine, and 1% NEAA without antibiotics. Cells were diluted to 5 × 10^4^ cells/mL and polybrene (EMD Millipore) was added to a final concentration of 10 μg/mL. Cells were transduced with different viral loads (ranging from 1 to 500 μL of viral supernatant per 1 mL of cells) to obtain cell populations with different multiplicities of infection (MOIs). After 3 days of viral transduction, cells were treated with puromycin (Invivogen) to select for cells that had successfully integrated viral DNA. C3H were treated with 5 μg/mL, C2C12 were treated with 5 μg/mL, and U2OS were treated with 1 μg/mL puromycin for up to 1 week. Cell populations were further expanded and grown for fractionation, Western blotting, and immunofluorescence.

### Western blotting

For Western blotting, cells were resuspended in 2X Laemmli buffer (4% SDS, 10% 2-mercaptoethanol, 20% glycerol, 0.004% bromophenol blue, and 0.125 M Tris-HCl pH 6.8) and boiled for 5 minutes. Samples were run on a Novex Tris-Glycine gel (Life Technologies) using FASTRun Buffer (Fisher Scientific). Samples were then transferred to a nitrocellulose membrane (Life Technologies). Membranes were rinsed twice with Tris-buffered saline (TBS) with 0.1% Tween-20 (Tw) and then blocked with 5% bovine serum albumin (BSA) in TBS/Tw. Membranes were incubated with primary antibody diluted in 0.5% BSA in TBS/Tw overnight at 4°C. Membranes were then washed 6 times with TBS/Tw and incubated with HRP conjugated secondary antibodies in TBS/Tw for 1 hour at room temperature. Signals were then developed using an enhanced chemiluminescence kit (Thermo Fisher) for 5 minutes before exposure to film.

### Immunofluorescence

For immunofluorescence staining, cells were plated on sterile glass coverslips and allowed to grow overnight. 24 hours after plating, cells were rinsed with Dulbecco’s phosphate buffered saline (DPBS with calcium and magnesium) and fixed using 2% paraformaldehyde (PFA) (Electron Microscopy Sciences) in PBS for 20 minutes. Samples were rinsed three times with phosphate buffered saline (PBS) and blocked for 15 minutes using PBS with 5% goat serum (Jackson ImmunoResearch Laboratories) and 0.5% Triton X-100 (Tx) (Fisher Scientific). Samples were then incubated with primary antibody diluted in PBS with 1% goat serum and 0.1% Tx overnight at 4°C. After washing with PBS/Tx (0.1%) 4 times, samples were incubated with Alexa Fluor conjugated secondary antibody diluted in PBS/Tx (0.1%) at room temperature for one hour. Samples were finally washed twice with PBS/Tx (0.1%), incubated with DAPI at room temperature for 10 minutes, and then washed twice with PBS and mounted on glass slides using Aqua-Poly Mount (Polysciences).

For digitonin permeabilization of C3H cells, 4 × 10^4^ cells were plated on sterile glass coverslips coated with 0.2% gelatin in a 24-well plate. 24 hours later, cells were fixed and treated with either 40 μg/mL or 1 mg/mL digitonin in PBS at room temperature for 5 minutes. Samples were then washed 3 times with PBS and blocked using PBS with 5% goat serum at room temperature for 15 minutes. Samples were incubated with primary antibody diluted in PBS with 0.5% goat serum overnight at 4°C. The next morning, samples were washed 4 times with PBS and incubated with secondary antibody in PBS for 1 hour at room temperature. Samples were then stained with DAPI, washed with PBS and mounted on glass slides using Aqua-Poly Mount.

### Light Microscopy and Quantification

Confocal images were acquired on a Zeiss 780 or a Zeiss 880 Airyscan laser-scanning confocal microscope with a 63X PlanApo 1.4 NA objective. Contrast adjustment of the representative images was performed with ZEN software (Zeiss). 10 or more images from each stably or transiently transduced cell population of the lowest expression levels were randomly chosen and the NE/ER ratio was quantified. Lamin A staining was used to outline the nucleus and the area of NE and ER were defined by -0.5 to 0 μm (NE) and +0.5 to +1 μm (ER) relative to the edge of the nucleus using the *“Enlarge”* function in ImageJ (NIH). Total fluorescent intensities of V5 staining in both areas were measured and normalized to the calnexin staining of the same area. The ratio of NE/ER was then calculated by dividing the normalized V5 signals in the NE to the normalized V5 in the ER.

The co-localization analysis was performed with the *“Coloc2”* function in ImageJ. Where necessary, raw images was processed using the rolling-ball *“background subtraction”* function in ImageJ. Control and test images were processed with identical parameters. Representative images were prepared with automatic Airyscan processing in ZEN.

## Supporting information

## Acknowledgements

LG and JRY acknowledge support from the National Institutes of Health Common Fund 4D Nucleome Program (Grant 1U01DA040709). Additional support was provided by NIH grants R01GM28521 to LG and P41GM103533 to JRY.

## Author Contributions

LG and JRY designed the project and supervised its execution; LG drafted the manuscript; LCC carried out the immunofluorescence analysis, fractionation of transduced cells, Western blotting and prepared figures and tables; SB executed the MudPIT analysis and aided data interpretation; CL developed the subcellular fractionation methods and prepared fractions for proteomics; LB did the molecular cloning and prepared stably transduced cell lines; SMB carried out data analysis and prepared tables; OT developed initial versions of the subcellular fractionation methods; XZ assisted with data interpretation

## Supplementary Figure Legends

**Figure S1.**
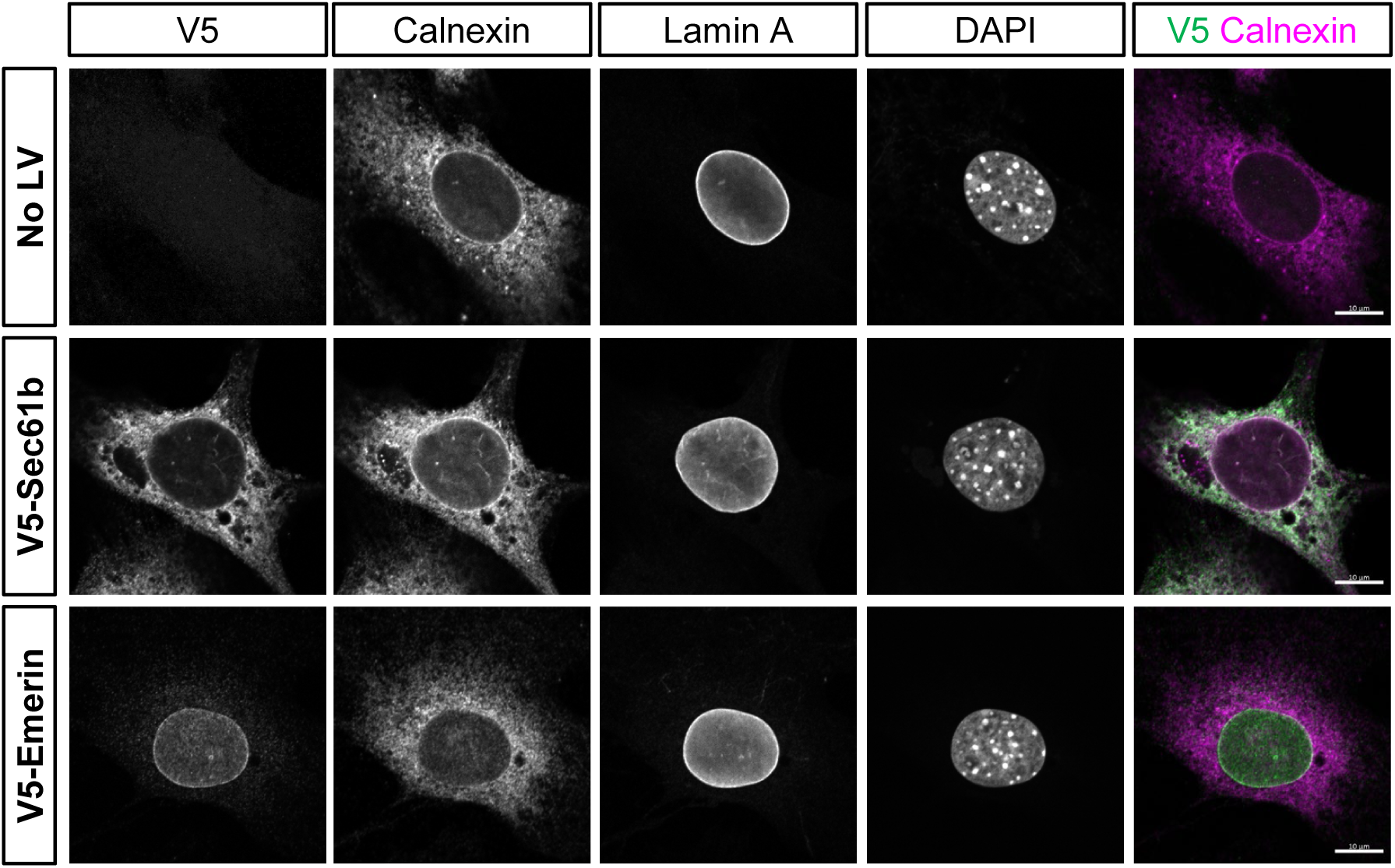
Immunofluorescence labeling of C3H cells stably tranduced with NE and ER control proteins. C3H cells were either treated with no lentivirus, or were stably transduced at low MOI with lentiviruses expressing either V5-Sec61b or V5-Lem2. Shown are representative immunofluorescence images of cells co-labeled with antibodies to the V5 epitope tag, calnexin, and lamins A. DNA staining (DAPI) and a merge of V5 and calnexin labeling is shown in the right panels. Scale bars, 10μm.

**Figure S2.**
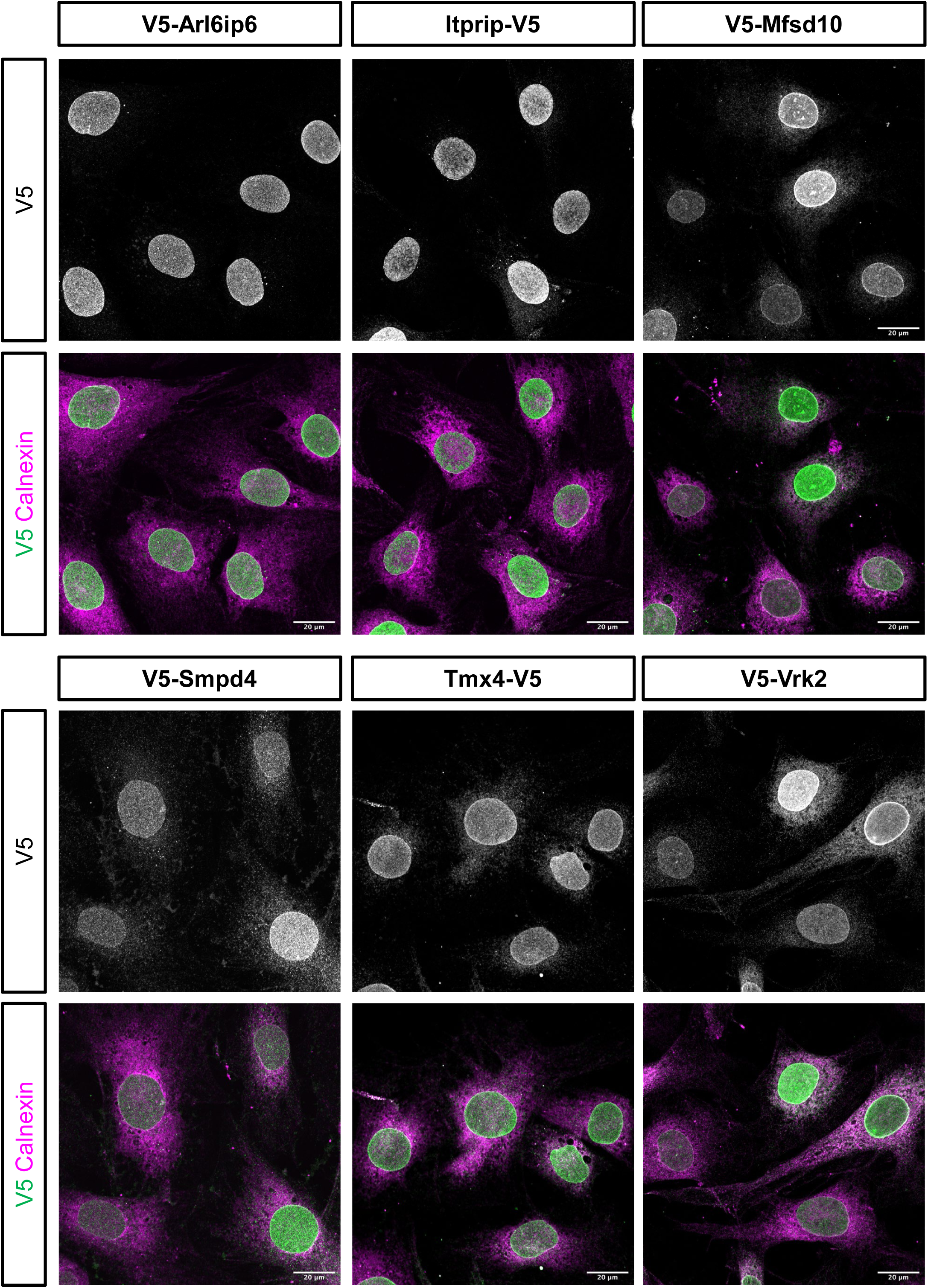
Immunofluorescence labeling of lower magnification fields of ectopically expressed targets. C3H cells were stably transduced with the indicated constructs (or in the case of Smpd4, transiently transduced) and were examined by immunofluorescence staining to detect the V5 tag and calnexin. Shown are representative lower magnification fields. Scale bars, 20μm.

**Figure S3.**
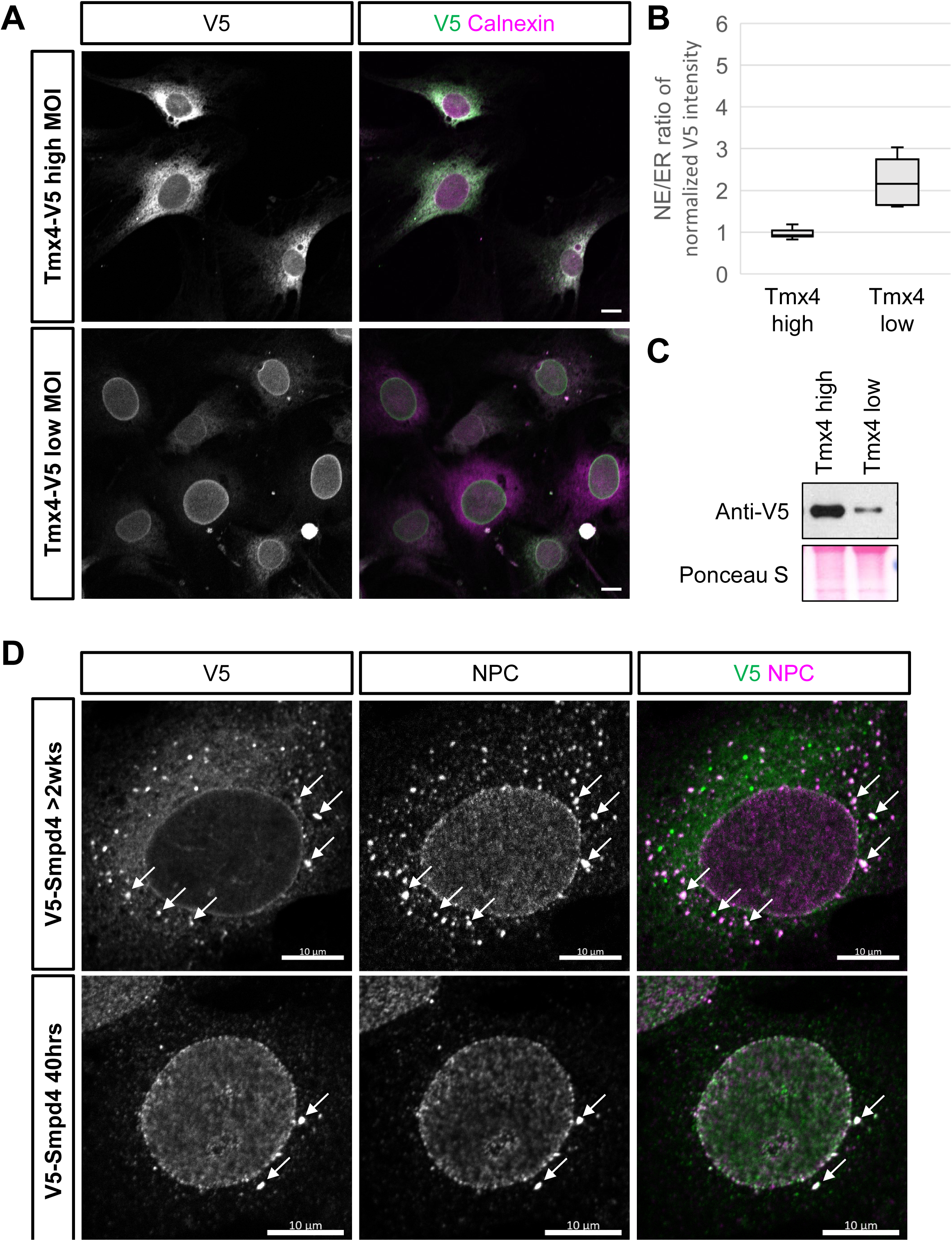
Comparison of immunofluorescence localization of Tmx4 and Smpd4 in cells expressing low vs high amount of ectopic protein. (A) C3H cells stably transduced with either low or high MOI lentiviruses expressing Tmx4-V5 were co-labeled with antibodies to the V5 epitope tag and calnexin. Shown are fluorescence images of V5 staining only (left) or V5 staining merged with calnexin labeling (right). Scale bars, 10μm. (B) Plot showing the NE/ER ratio of Tmx4, calculated as in Fig. 4B, in low vs high MOI populations. (C) Western blot showing level of ectopic Tmx4 expression in low vs high MOI populations, with corresponding Ponceau staining. (D) Immunofluorescence images showing C3H cells examined either shortly after transduction with a lentivirus expressing V5-Smpd4 (40 hours), or selected for stable lentiviral expression (>2 weeks). Cells were co-labeled with anti-V5 and the RL1 antibody recognizing FG repeat Nups of the NPC. Small numbers of cytoplasmic foci positive for both Smpd4 and RL1 (arrows) are seen in the 40 hour cells. The number of these cytoplasmic foci is greatly increased in in the >2 week transduced cells. Not all V5-labeled cytoplasmic foci are strongly positive for RL1 antigens. Scale bars, 10μm.

**Figure S4.**
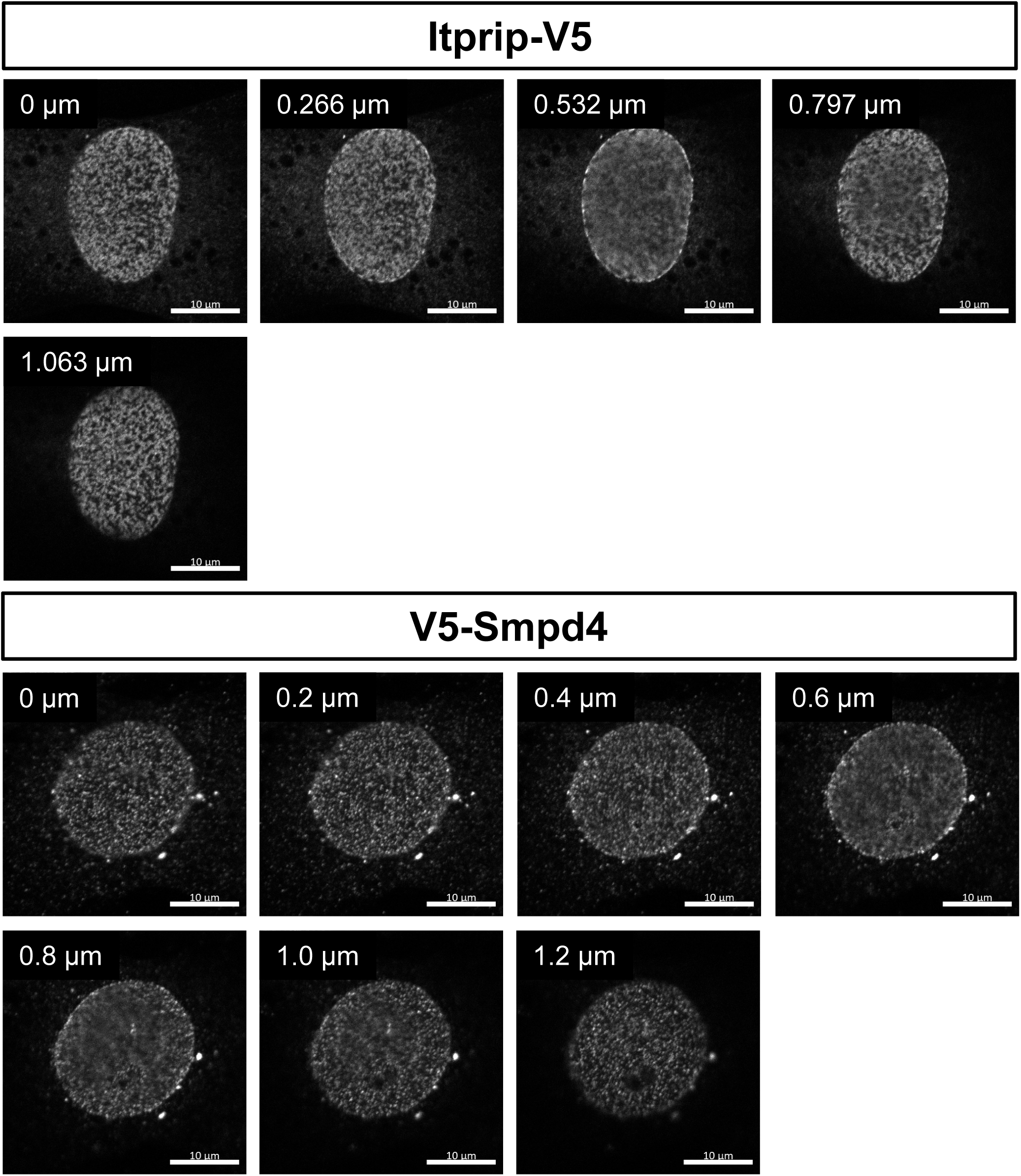
Z-stacks of confocal images of C3H cells expressing V5-tagged Itprip or Smpd4. The Z-position of each image is indicated. Note that C3H nuclei are very flat in the Z dimension. Scale bars, 10μm.

**Figure S5.**
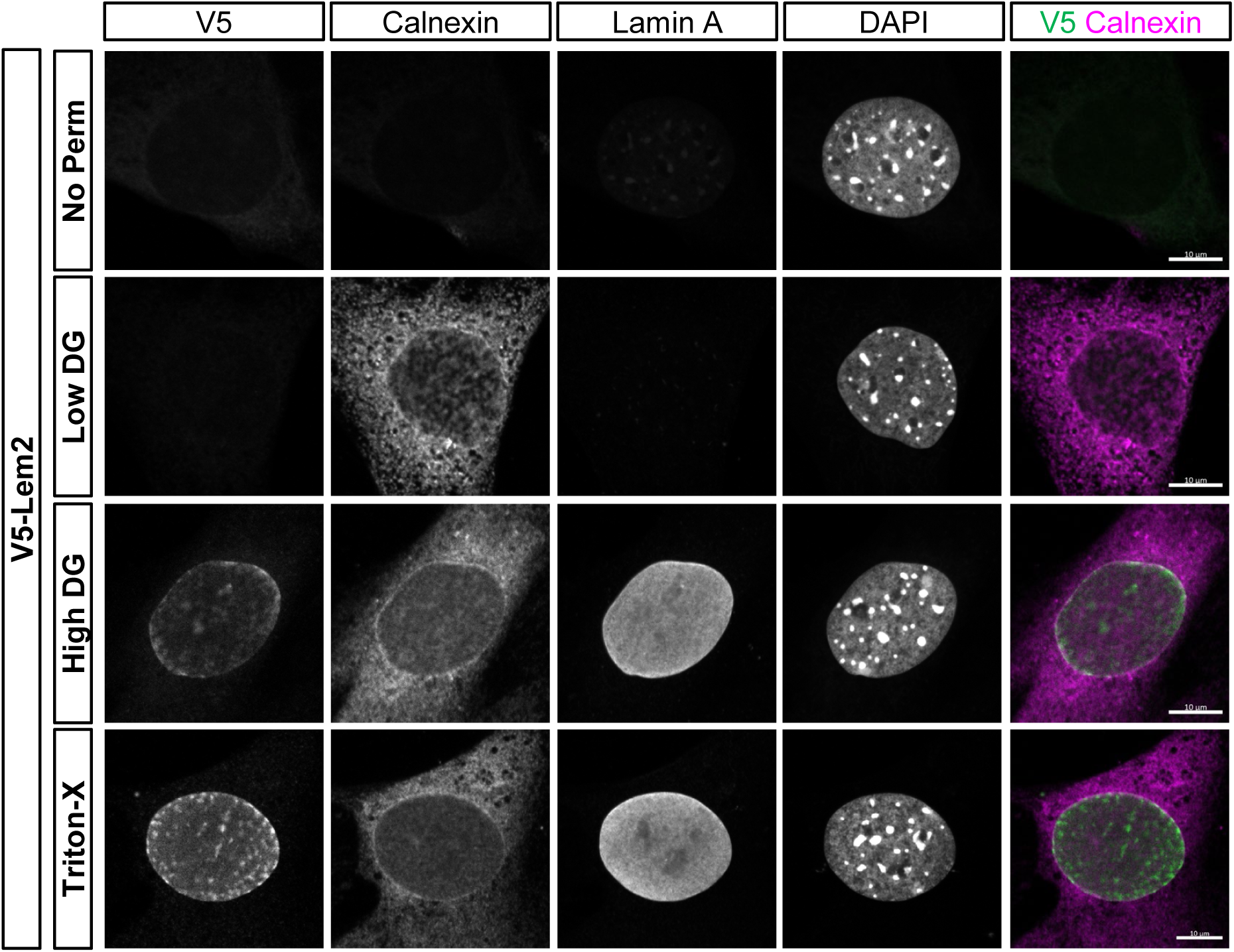
Accessibility of ectopically expressed V5-Lem2 to antibodies after permeabilization with low vs high digitonin. Shown are immunofluorescence images of C3H cells incubated with a mixture of antibodies to V5, calnexin, lamin A and DAPI after no permeabilization, or permeabilization with low or high digitonin, or with Triton X-100 as indicated. Scale bars, 10μm.

**Figure S6.**
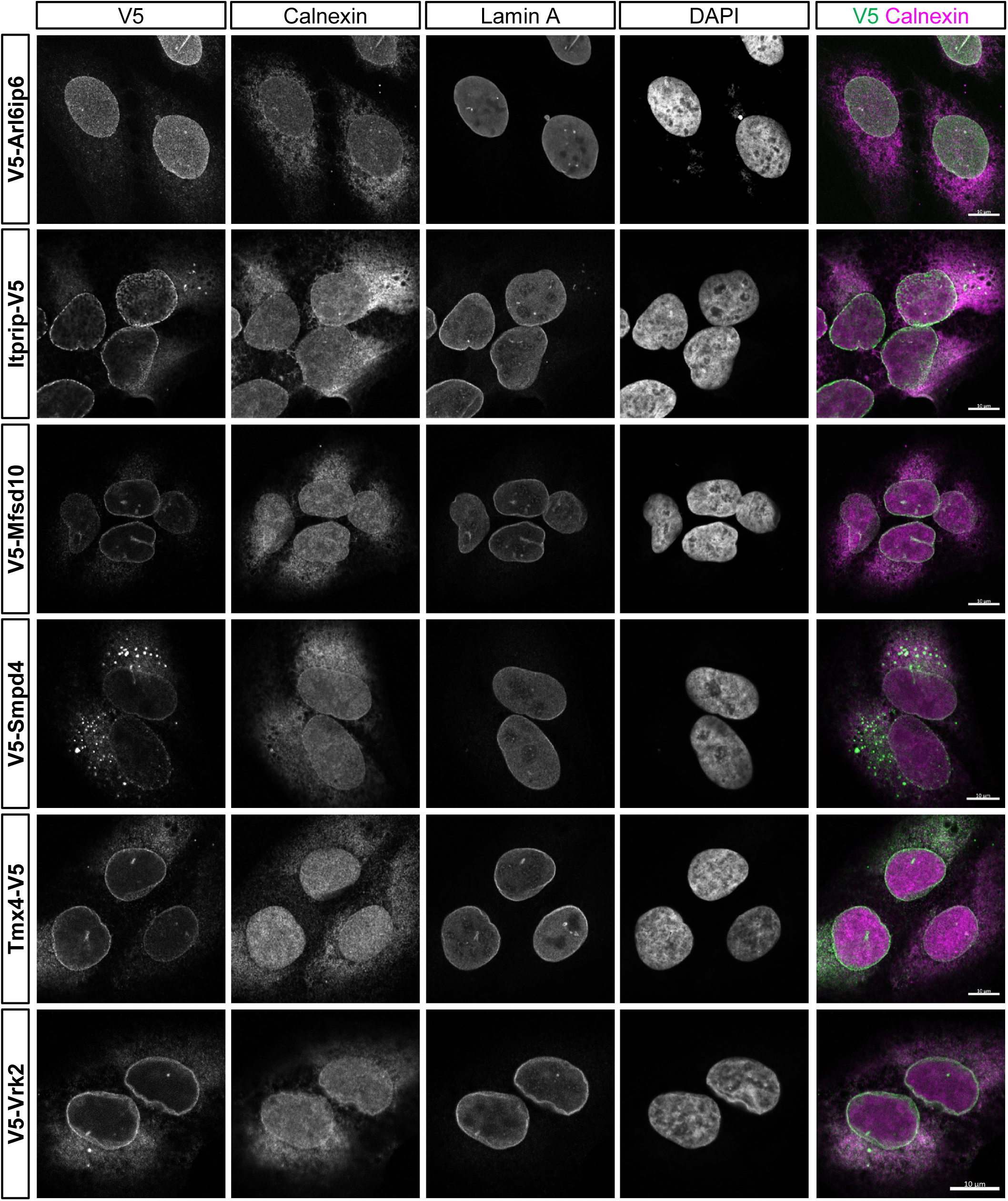
Targeting of ectopically expressed target proteins to the NE in U2OS cells. Cells were stably transduced with lentiviruses expressing the constructs indicated. Shown are representative immunofluorescence images, with antibody staining and representation as in Fig. S1. Scale bars, 10μm.

**Figure S7.**
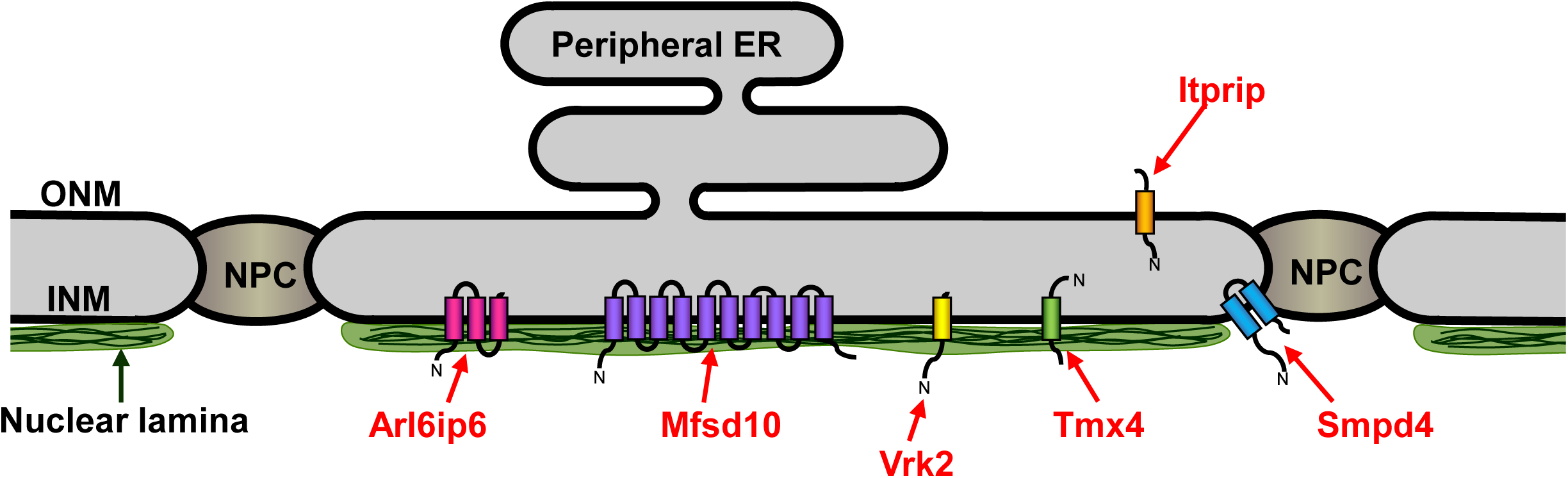
Diagram depicting the localizations of the proteins analyzed in this study deduced from IF microscopy and digitonin permeabilization of C3H cells.

**Supplementary Table S4.**
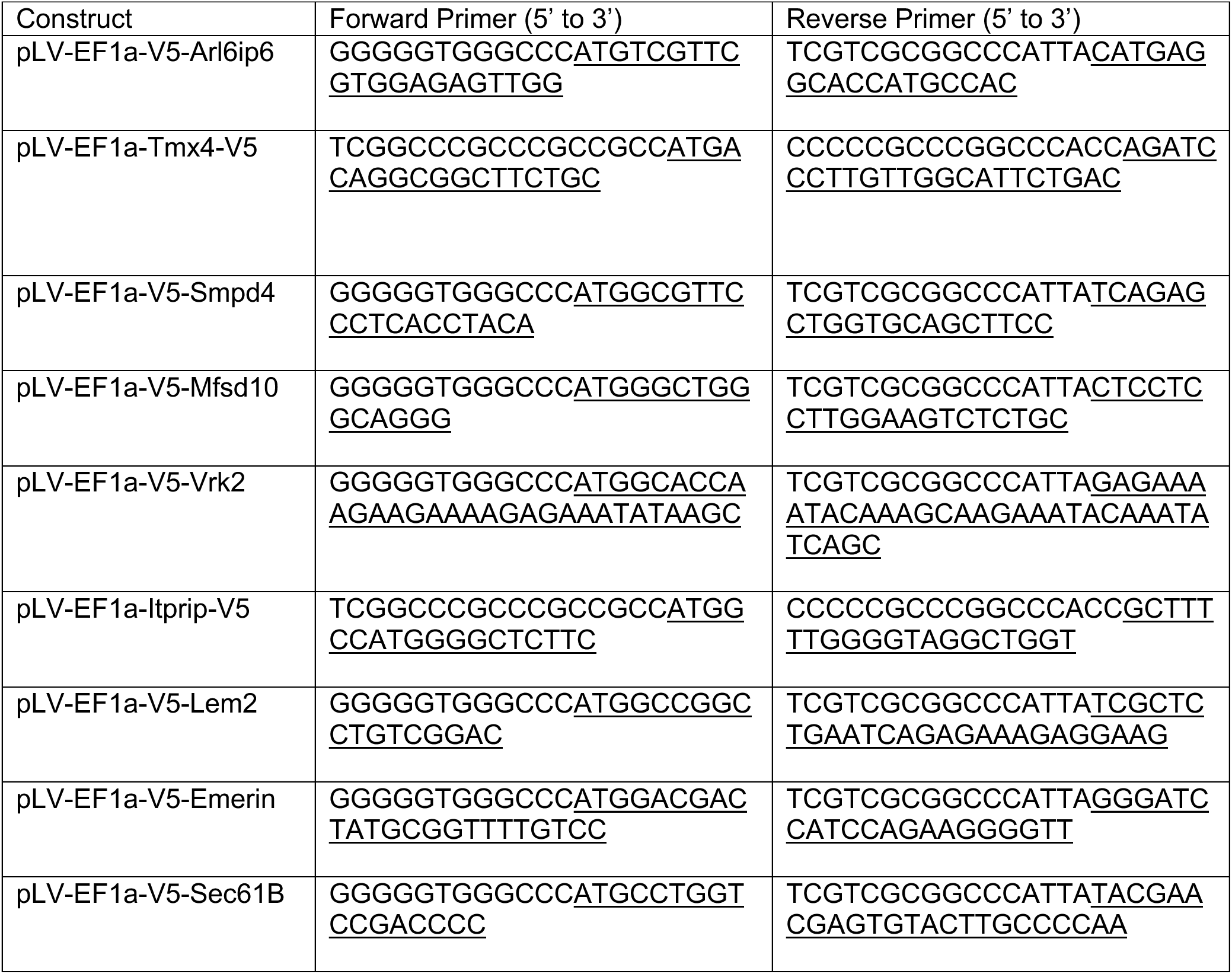
Primers used in this study.

